# Individual differences in perceptual decision making reflect neural variability in medial frontal cortex

**DOI:** 10.1101/139493

**Authors:** Tomoki Kurikawa, Takashi Handa, Tomoki Fukai

## Abstract

Decision making obeys common neural mechanisms, but there is considerable variability in individuals’ decision making behavior particularly under uncertainty. How individual differences arise within common decision making brain systems is not known. Here, we explored this question in the medial frontal cortex (MFC) of rats performing a sensory-guided choice task. When rats trained on familiar stimuli were exposed to unfamiliar stimuli, choice responses varied significantly across individuals. We examined how variability in MFC neural processing could mediate this individual difference and constructed a network model to replicate this. Our model suggested that susceptibility of neural trajectories is a crucial determinant of the observed choice variability. The model predicted that trial-by-trial variability of trajectories are correlated with the susceptibility, and hence also correlated with the individual difference. This prediction was confirmed by experiment. Thus, our results suggest that variability in neural dynamics in MFC networks underlies individual differences in decision making.

## Introduction

Animals need to respond to sensory input from the environment and make adaptive decisions, even when the sensory information is ambiguous. When animals are familiar with given sensory stimulus, their behavioral responses are stereotyped. However, when sensory input is unfamiliar to animals, their responses are more variable and exhibit a highly probabilistic nature. Decision making in such ambiguous situations is expected to show a wide spectrum of individual differences depending on the behavioral traits of each subject.

Many studies have explored the general tendencies and underlying neural mechanisms of probabilistic decision making [1–3]. In particular, many studies have suggested the crucial role of choice-specific neural activity sequences in decision making behavior [4–6]. However, surprisingly little is known about the characteristic features of neural activity that influence the behavioral variability of individual animals. In this study, we experimentally and computationally explore such features and the underlying mechanisms in choice-specific sequences in medial frontal cortex (MFC). Inactivation of the MFC is known to impair motor responses driven by sensory input [7], and recent studies have shown that the MFC is engaged in decision making and goal-directed behavior [8].

Choice-specific neural trajectories have been found in the MFC of rats performing an alternative choice task in response to auditory stimuli [9]. In this task, after rats had been trained with familiar tone stimuli, they were required to respond to unfamiliar tone stimuli that had not been used during training. MFC neurons in these animals formed choice-specific trajectories for familiar stimuli. Furthermore, the probabilistic choice responses of the rats showed substantial individual differences in psychometric curves, presumably reflecting different preferences and/or strategies in decision making. These findings motivated us to explore the cortical mechanisms underlying the reward-driven formation of choice specific sequences and their influences on the individual differences. Cortical mechanisms have been extensively studied to account for general tendencies in decision making behavior [10, 2, 3], but little has been explored about the neural correlates of individual differences.

In this study, we first address how neural trajectories emerge for familiar stimuli and guide decision making for unfamiliar stimuli. We then ask whether and how the properties of these trajectories influence choice responses and the resultant psychometric curves. For this purpose, we construct a recurrent network model [11–14] based on experimental observations from the rat MFC, and train it with a reinforcement learning algorithm for an association task [15–17]. Our model generates a spectrum of decision behavior that is compatible to the observed variability in behavioral performance across different rats. We found that susceptibility of neural trajectories to perturbation predicted variability in behavior and that trial-by-trial variability reflected the susceptibility and, consequently, the behavioral variability. Our results suggest that the susceptibility of neural dynamics correlate the trial-by-trial variability in MFC with individual differences in choice responses to uncertain stimuli.

## Results

### Behavioral variability in decision making with unfamiliar sensory cues

Our hypothesis is that individual differences in ambiguous choice responses partly emerge from neural dynamics formed in MFC through previous experiences. We examined this hypothesis in a sensory-guided alternative choice task. Rats were trained to make either Left or Right licking in response to high-frequency (13 kHz) or low-frequency (10 kHz) auditory cues (called familiar cues), respectively (Figure 1A). Each cue was presented for 200 ms, and correct choices after the cue presentation were rewarded. Among 36 trained rats, 21 rats reached a criterion correct rate (75 %), and multi-neuron recordings were performed from MFC of these rats. During the recordings, the rats were exposed to unfamiliar cue tones (10.5 to 12.5 kHz) in 20% of trials besides the familiar tones. Because these tones were unfamiliar to the rats, their choice responses were ambiguous and probabilistic. Finally, eight rats yielded sufficiently many neurons for the present analysis. Further experimental details are found in Methods.

**Figure 1:**
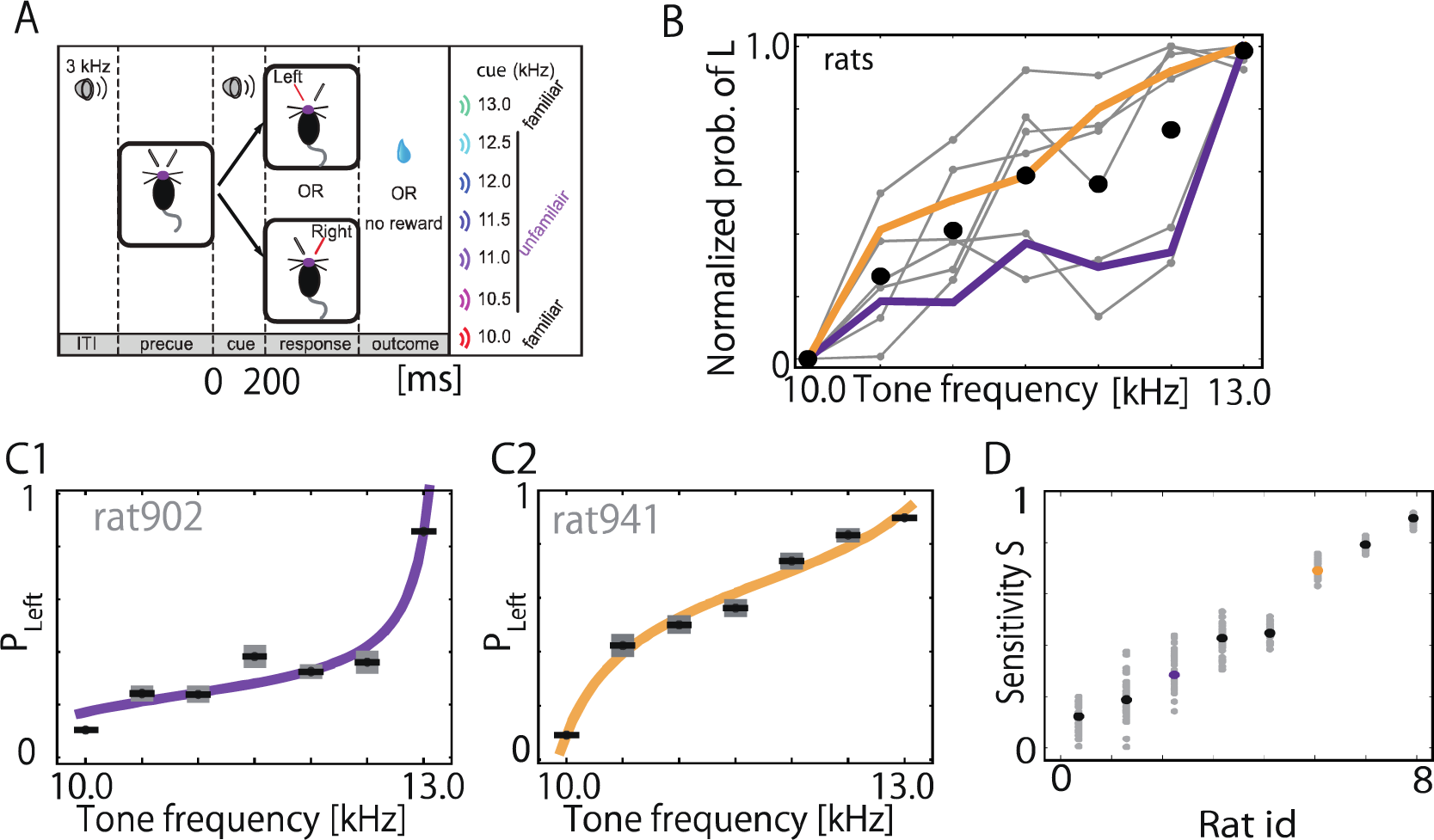
Individual differences in probabilistic choices across rats. Psychometric curves and task-related population neural activity were studied in eight rats. A, Schematic illustration of sensory-guided decision making task. B, Normalized psychometric curves (gray) significantly varied across rats (*N* = 8). In each rat, the probability of left licking was normalized such that *P*(10 kHz)=0 and *P*( 13 kHz)=1. Orange and purple curves are typical examples with high and low sensitivity to auditory cues, respectively. Filled circles indicate the average Left choice probabilities over the eight rats. C, The typical psychometric curves of low and high sensitive rats were fitted with tangent functions. Error bars show standard deviations of resampled ensembles. D, Sensitivity *S* for all rats is plotted. 30 samples of psychometric curves were generated by resampling for each rat and the sensitivity values S of the resampled (gray) and original (purple for the insensitive rat and orange for the sensitive rat) curves were calculated. The rats are sorted in the abscissa in an increasing order of their original sensitivity values.

This experiment revealed an interesting feature of decision-making in unfamiliar situations: The choice behavior of the rats showed large differences across the rats. The choice probability of Left licking generally increased with the tone frequency in all rats, but psychometric curves for unfamiliar cues exhibited large individual differences (Figure 1B). Some rats varied choice probability sensitively to the frequency of unfamiliar cues (Figure 1C2), but other rats made biased or chance-level choices irrespective of the tone frequency (Figure 1C1). Throughout this study, we quantified the individual differences in psychometric curves by fitting these curves with a nonlinear function *P*_L_ (Methods). Choice probabilities are inevitably deflected by sampled data. Therefore, we examined the stability of parameter fitting as follows. We resampled a different set of 30 trials out of the entire data set (comprising several tens of unfamiliar trials and several hundreds of familiar trials) for each tone and rat. This allows us to generate a psychometric curve per rat. Then, for each rat we repeated this procedure 30 times to generate 30 samples of psychometric curves and to collect the corresponding 30 values of sensitivity *S*. We plotted these values in an increasing order of the original sensitivity of the rats (Figure 1D). Though the variance was large in some rats, the average values of *S* monotonically increased while preserving the serial order of *S*, indicating that the sensitivity adequately characterizes the behavioral tendency of each rat irrespective of data sampling. We further confirmed the stability of our fitting scheme by choosing a different fitting function (Figure S1).

### Reinforcement learning for reservoir network model

To clarify how the behavioral differences may emerge among rats, we constructed a stochastic recurrent network of spiking neurons based on the anatomical structure from MFC (Figure 2A). Our model extends the reservoir computing proposed for supervised motor learning [11–14] to reward-driven sequence learning [15–17]. The recurrent network (reservoir) consists of 5000 excitatory and 1000 inhibitory neurons, and a subset of these neurons receive random projections from 200 input terminal neurons having different preferred tone ‘frequencies’. Anatomically, the input terminal and recurrent network may correspond to the auditory cortex and MFC, respectively [18, 19]. Random projections were assumed because tone-selective MFC neurons showed no systematic frequency dependence [9]. Excitatory neurons in the reservoir project to two rate-coding neurons (readout neurons), L-neuron and R-neuron, which are mutually inhibiting. The readout neurons send feedback connections to the reservoir. We also assumed these readout neurons are representative of motor cortex generating licking behavior [20]. The network model makes a decision when the difference in output between L and R neurons reaches a threshold *θ*. Due to the noise applied to the reservoir and trial-to-trial variability in the initial states, the decisions are stochastic even for an identical sensory input.

**Figure 2:**
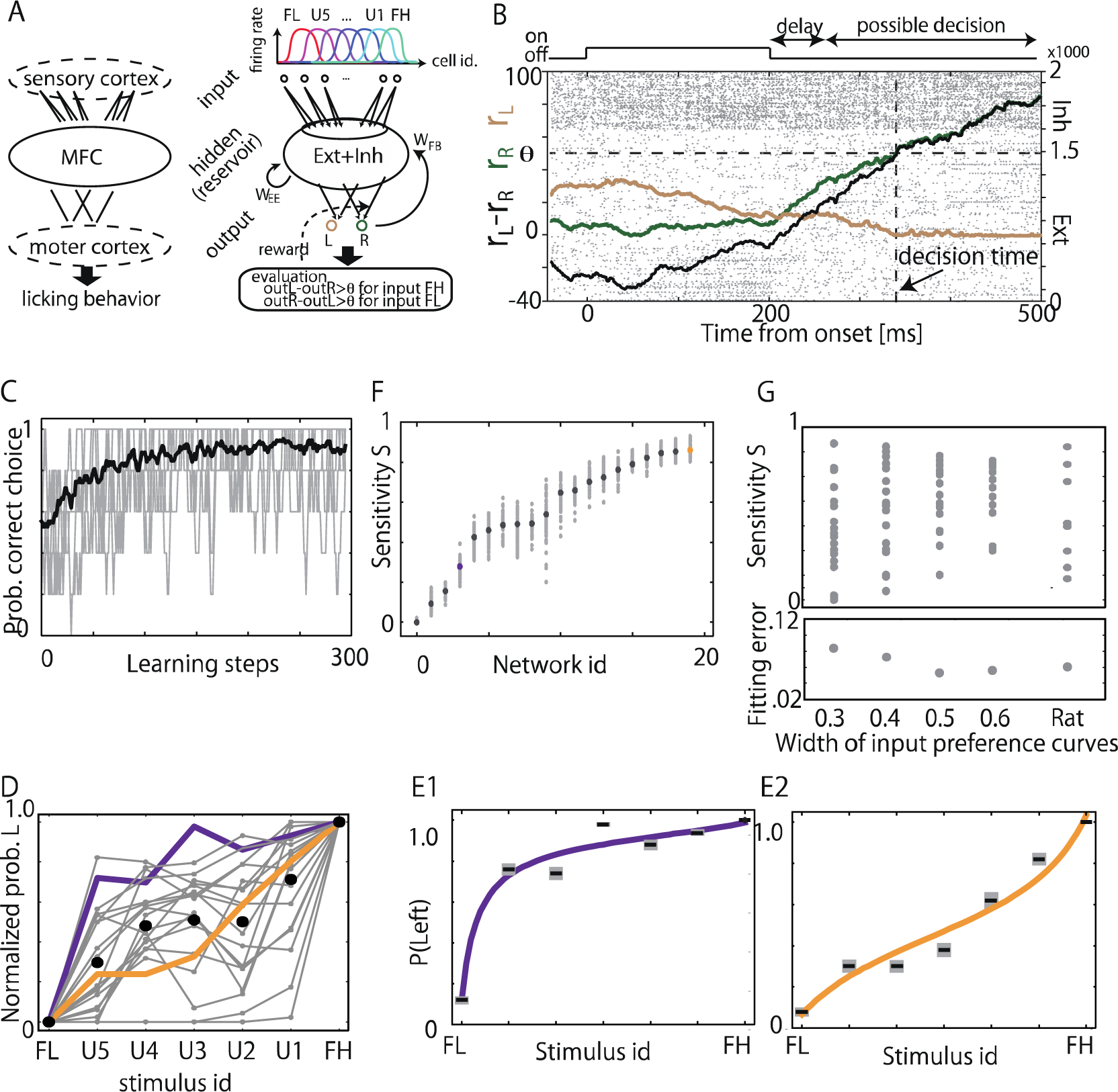
Responses of network models reproduce individual difference. A, Schematic illustrations of the presumable anatomical circuits engaged in the present task (left) and the model network structure (right) are shown. B, Spike raster of the reservoir (dots) and the firing rates of L (rL, gold) and R (rR, green) neurons are shown for a single trial. The figure shows randomly sampled 500 excitatory neurons (ENs) receiving the input projections, 1000 ENs receiving no input projections, and 500 inhibitory neurons. C, Learning performance of the model. The probability of correct choices was evaluated every 5 learning steps. Black and gray lines indicate the average performance over 30 different realizations of the network model and the performance of five examples among them, respectively. D, Psychometric curves of 20 successful learners (gray) and two typical curves with high (orange) and low (purple) sensitivity are plotted. E, The typical curves are well fitted with tangent functions. F, Sensitivity S of the original (black) and resampled (gray) psychometric curves for 20 successful learners are plotted. Orange and purple circles indicate 5 for the original psychometric curves for sensitive and insensitive networks. Panels D-F are compared with Figures 1B-D. G, Comparison of distributions of 5 in models and rats. Upper figure shows 5 for each fitting curve against width of preference curves. We plotted 5 for 18, 20, 16 and 14 networks for the width = 0.3, 0.4, 0.5 and 0.6, respectively, as well as those for eight rats. Bottom, fitting errors averaged over networks and rats are shown.

We trained the network model through a reward-driven learning rule such that it correctly generated “Left” or “Right” choice for familiar “high-frequency” (FH) or “low-frequency” (FL) input, respectively (Methods). To take advantage of sequence generation [21, 22], we made recurrent excitatory connections obey a lognormal weight distribution, which has been ubiquitously found in cerebral cortices [22, 23]. We also assumed a minimal model in which only readout connections were modifiable. Although recurrent connections were unchanged to keep the lognormal weight distribution stable against spiking dynamics, the reservoir dynamics is modulated through the feedback connection from the read-out neurons. We then introduced the following reinforcement learning by using eligibility trace [15–17]:

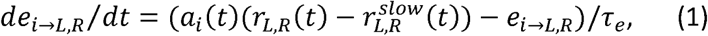

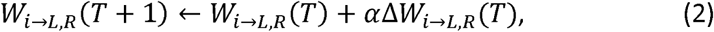

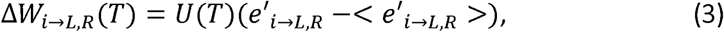

where 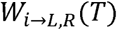 is the weights of readout connections from the *i*-th reservoir neuron to L- or R- neuron at learning step *T*, *α* is a learning rate, *α_i_*(*t*) is the average activity of presynaptic reservoir neuron (Equation 13 in Methods), and *r_L,R_*(*t*) and 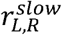 are the firing rate of each readout neuron and its low-pass-filtered version, respectively. Throughout the paper, *α* = 10. Eligibility trace 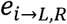 is assigned to each readout connection and measures the extent to which a particular connection contributes to decision making at a particular learning step. In Equation 1, the eligibility trace is calculated from correlations between *a_i_*(*t*) and high-pass-filtered readout activity 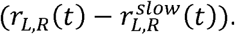. In Equation 3, 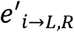 is the normalized eligibility trace of this connection, and 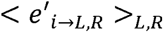 is the average over all such connections (Methods). Modification of the readout connections (Equation 2) occurs only after one learning step finishes. After a success trial (learning step) *U* is taken to be positive (i.e., potentiation of influential connections), or *U* is negative after a failure trial (depression of such connections). In each trial, the network was exposed to one of the familiar inputs from 0 to 200 ms and made a decision within 250 to 900 ms after the stimulus onset. If the readout neurons did not reach the threshold within this interval, the trial was failed and a new trial was started. Though the learning rule is somewhat heuristic, the minimal model accounted for most of the essential results of experiment as shown below.

Figure 2B shows a typical example of a rewarded trial during learning. We trained 30 networks with different realizations of recurrent connections obeying an identical weight distribution (Methods). For each network, either FH or FL is randomly applied during learning. After 300 training steps, more than the two-third of model networks (20 networks) achieved a criteria correct rate of 75 % (Figure 2C). Successful learners occupied similar fractions in both models and rats, indicating that the task difficulty was adequately modeled. Throughout this study, we analyzed the neural dynamics of these 20 successful learners and 8 rats, and compared them in their behavioral variabilities.

Next, we analyzed the highly stochastic choice responses of our models to unfamiliar cues (U1 to U5 in Figure 2A). As in the rats, the psychometric curves of the successful learners displayed large individual differences (Figure 2D), where we used the same fitting function (Figure 2E) as used in the rats (Figure 1C). The sensitivity values thus obtained for the different models were robust against the resampling of psychometric curves (Figure 2F). As the width of tuning curves (see Figure 2A) influences the initial network states set by external stimuli, it also influences the behavioral characteristics of each network. In Figure 2G, we compared the distributions of sensitivity values of the models for different widths of tuning curve. The model networks tend to exhibit a broader sensitivity distribution for a narrower width (i.e., a higher input selectivity). When the width is 0.3 to 0.4, the ranges of sensitivity covered by the models are as broad as the range covered by the rats. Below, we use the width of 0.4 as it gives a smaller average fitting error than the other value.

### Neural trajectories formed in MFC and the reservoir

We have shown that the network models produce the divergent psychometric curves that are consistent with those observed in experiment. However, whether neural activities in the reservoir and in the MFC are also similar has yet to be clarified. To this end, we first studied the time evolution of neural activities. Figures 3A and 3B compare neural activities averaged over familiar trials between in MFC and in the reservoir under given cue and choice conditions. Both experimental and modeled neural activities exhibited distinct trajectories when neurons were sorted according to their times of peak firing rates in correct trials for given stimulus. However, neural activities sorted in the peak-time orders observed in different stimulus conditions showed no clear trajectories. These results demonstrate that the neural trajectories are choice-specific in both MFC and the reservoir.

**Figure 3:**
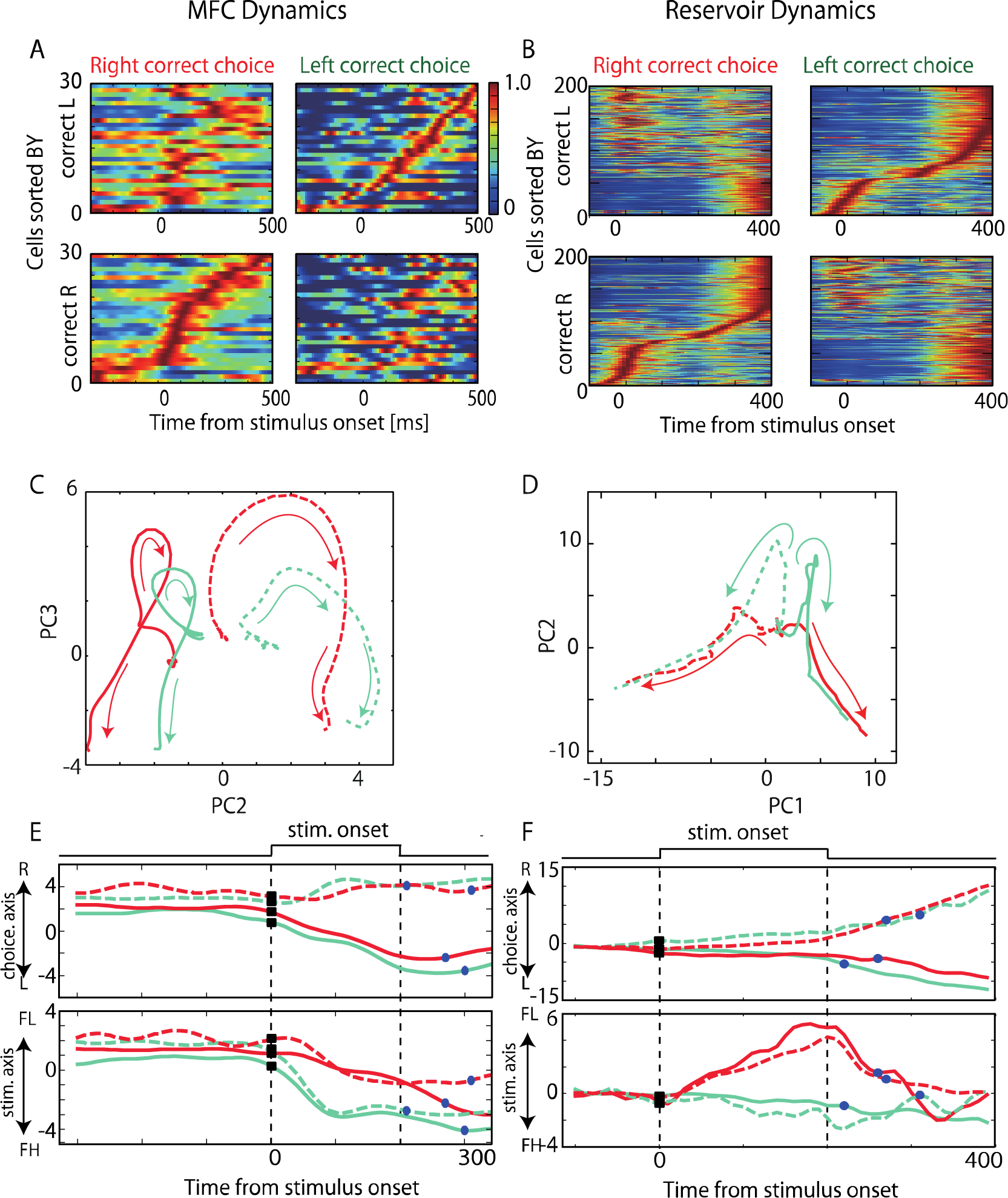
Neural dynamics in MFC and in the reservoir. A, Neural activities of 30 most active neurons in a rat out of 44 were averaged and normalized over success trials with FL (left) and FH (right) tones. Neural activities were sorted in the temporal order of peak responses to FH (top panels) or FL (bottom panels). B, Neural activities in a model are shown in a similar fashion to A. 200 most active excitatory neurons out of 5000 are used. C and D, Neural trajectories in the MFC (in C) and the reservoir (in D) are projected onto a two-dimerisiorial PC-space. We averaged neural activities over 30 neurons showing the largest difference between firing rats before and after familiar stimuli in the MFC and 100 excitatory neurons showing the highest firing rate in the reservoir under each condition of stimuli and choices. The second and third PCs were used because PCI merely represented the presence of cues without tone specificity. E, Time evolution of population neural activity in a rat was projected onto the choice (top) and stimulus axes (bottom) for FH (light green) and FL (red). Solid and dotted lines show neural trajectories for Left and Right choices, respectively. Squares and circles indicate stimulus onset and mean RT. F, Time evolution in a model is shown in a similar fashion to C.

We analyzed the principal components (PC) of population neural responses in MFC and the reservoir in Figures 3C and 3D, respectively. In MFC, decision making with familiar cues formed neural trajectories selective to stimulus and choice. Note that the first principal component (PC1) represented a frequency-nonselective component of auditory responses (Figure S2) and hence is not shown here. In the reservoir, similar condition-selective trajectories are formed except that PC1 and PC2 in the former corresponded to PC2 and PC3 in the latter, respectively. Frequency-nonspecific stimulus-evoked responses (PC1 in the experiment) were almost missing in our models.

We then investigated how these familiar trajectories lead to correct choice responses in MFC and the reservoir networks. As PCA does not identify task-relevant axes, we applied a linear regression method (Mante et al., 2013 and Methods) to trial-averaged neural activities for given combination of stimulus and choice, and identified two axes: the stimulus axis explains the maximal difference in trajectories between FH and FL while the choice axis reveals the maximal difference between Left and Right choices. In both rats (Figure 3E) and models (Figure 3F), familiar trajectories projected onto the stimulus axis or the choice axis started to separate into correct choices almost simultaneously with cue onset. These results suggest that the integration of sensory evidence occurs during the cue presentation in both MFC and the reservoir.

The familiar trajectories also revealed interesting differences between the computational models and rats. First, the trajectory separation along the stimulus axis was generally smaller in the rats compared to the models. Though the cause of this discrepancy is not entirely clear, it was partly due to that the number of tone-selective MFC neurons was small and difference in their responses to different tones were also weak (at most several spikes per second in average). Second, trajectory separation along the choice axis stopped in the rats after the termination of sensory cues, whereas the separation continued to grow in the models presumably due to positive feedback from readout neurons. It was previously shown in the rat MFC that trajectories reach an almost maximal separation just prior to cue termination (Handa et al., 2017). The gamma-band power of the local field potentials also abruptly increased in MFC, indicating that some internal event, for instance the formation and transmission of a preparatory signal for choice response, might occur at this timing. Such an internal process was not modeled here, which might cause the discrepancy between rats and models. Except for these differences, our computational model well reproduced neural dynamics in the rat MFC.

### Neural dynamics in MFC and reservoir predicts choice behaviors

To confirm the importance of neural trajectories for choice behavior, we examined whether neural trajectories in MFC and our model have sufficient information about consequent choice responses. We predicted a probable choice from neural trajectories by using Fisher discriminant analysis (FDA) with a hyperplane that optimally divides the population activity patterns corresponding to Left and Right choices (Methods). The optimal hyperplane was determined from neural ensembles at the time of choice response for familiar trials, and we denote its normal vector as ***W***_opt_. In our model, if the readout neurons can discriminate optimally between Left- and Right-choice familiar trajectories, the normalized weight vector of readout connections should be close to ***W***_opt_. However, the two vectors cannot be exactly the same because the readout neurons are not passive linear filters but are active nonlinear filters performing temporal integration of inputs under mutual inhibition.

Averaged familiar trajectories projected onto ***W***_opt_ are shown for the rats (Figure 4A) and models (Figure 4B) for different cue and choice conditions. The projected trajectories linking given sensory cue to different choices were distinctive in both rats and models. Importantly, ***W***_opt_ obtained from familiar cues was also valid for discriminating between Left- and Right-choice trajectories evoked by unfamiliar cues. This means that unfamiliar trajectories converge at similar locations at which familiar trajectories also converge. We computed the distributions of projected activities in a rat (Figure 4C1) and a model (Figure 4C2) at the choice time points of Left-choice and Right-choice trials. In the rats, the activity distributions for different choices have small overlaps corresponding to incorrect choices. To infer trial-by-trial choice responses from the projected neural activity, we adopted a naive criterion boundary given as the mid-point of the means of Left- and Right-choice distributions. Then, Left-choice probability is predicted to be the area of the distribution lying on the left side of this boundary, and so on. On average about 75 % of neural trajectories in MFC were discriminable by this criterion, although the discriminability fluctuated across rats (Figure 4D). In the models, choice responses were correctly inferred in about 90 % of the trials, and fluctuations in the discriminability were small (Figure 4E). Thus, neural trajectories in MFC and the reservoir determine choice responses on a single-trail basis. This is expected in the model as it was trained as such, but the finding is non-trivial in the rat MFC.

**Figure 4:**
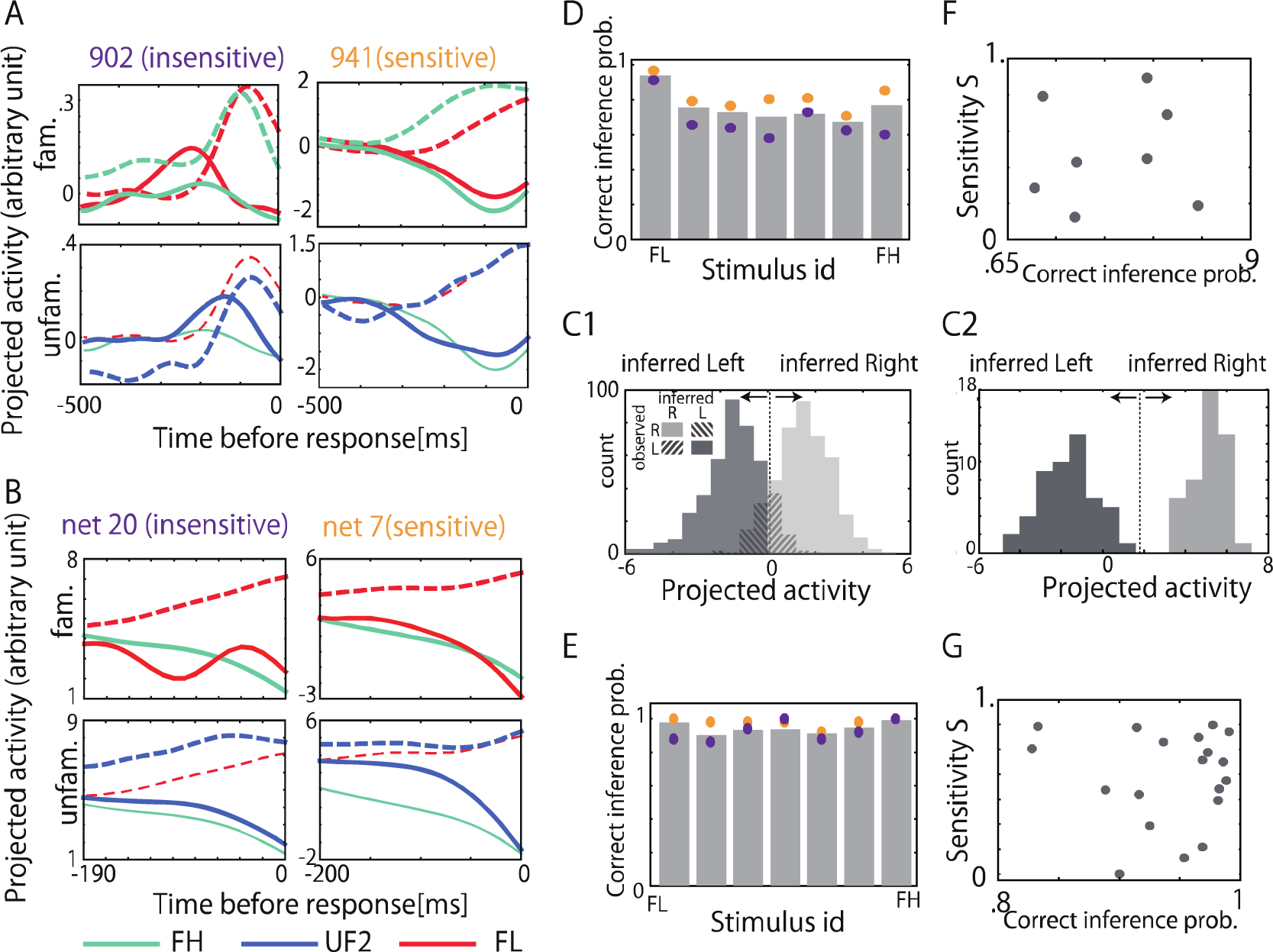
Predictions of choice behavior by MFC and reservoir dynamics. A and B, Neural trajectories were optimally separated by FDA in rats and models with different sensitivity, respectively. In both rats and models, projected trajectories are shown in Left- (solid) and Right-choice (dotted) trails for both familiar (top, FL and FH) and unfamiliar stimuli (bottom, U2). C1 and C2, Distributions of projected population activity are shown at the times of Left choices for FH (dark gray) and Right choices for FL (light gray) in the sensitive rat and sensitive net model shown in C1 and C2, respectively. Dotted lines show the midpoints between the mean values of the Left-choice and Right-choice distributions, and shaded area indicates the failure trials inferred from FDA. D and E, The probabilities that Left and Right choices are correctly inferred from the projected trajectories are shown in the rats and models, respectively. Gray bars show the probabilities averaged over all rats and all models, while orange and purple circles indicate the probabilities for sensitive and insensitive rats (D) or such models (E). F and G, Relationships between the probability of correct inference and sensitivity are shown for eight rats and twenty model networks, respectively.

However, the linear discrimination analysis was insufficient for predicting individual differences in choice responses across different rats and different models. We did not find significant correlations between the sensitivity to stimuli and the discriminability of neural trajectories in both rats (Figure 4F, p=0.86) and models (Figure 4G, p=0.82) although this could be partly due to small data size in the case of rats. These results suggest that the significantly different behavioral characteristics are not mere reflections of neural states at the choice point. Therefore, we investigate how neural population dynamics preceding the choice point determines the behavioral characteristics.

### Susceptibility of neural dynamics influences behavioral characteristics

Decision making in our model depends on the interplay between its internal dynamics and sensory input. Therefore, we expected that different characteristics of internal dynamics result in different behavioral characteristics. To characterize internal dynamics, we applied a perturbative input to networks receiving no sensory stimuli. The perturbative input was applied 30 times to each network at the identical neural state (perturbed state) to which sensory stimuli were applied in the previous simulations, and the Euclidian distances (susceptibility *χ*) among the perturbed trajectories and a non-perturbed trajectory were measured at 300 ms after the perturbation. The time of measurement did not change the essential results. Note that *χ* is defined for every different state in the neural state space (Methods). Perturbed neural trajectories evolving from a state with high *χ* may diverge broadly, while those trajectories from a state with low *χ* may hardly diverge.

We first show neural responses of a sensitive network and an insensitive network, respectively. For a particular state in the sensitive network, unfamiliar stimuli U1, U2 and U3 (close to FH) evoked trajectories evolving into Left choice, whereas those evoked by U4 and U5 (close to FL) evolved into Right choice, implying that the similarity of unfamiliar stimuli to familiar stimuli determined choice responses (Figure 5A1). In contrast, all unfamiliar trajectories resulted in Left choice for another state in the insensitive network, implying that internal dynamics governed the choice responses (Figure 5A2). We then show how perturbed and non-perturbed trajectories evolve in these networks from the same initial states. In the sensitive network, perturbed trajectories diverged broadly, implying that a weak perturbation tends to give high *χ* values (0.014, Figure 5B1). In contrast, the insensitive network generally yielded more localized perturbed trajectories and hence low *χ* values (0.0045, Figure 5B2). Thus, networks with high susceptibility responded differently to different unfamiliar stimuli, whereas those with low susceptibility responded similarly to different stimuli.

**Figure 5:**
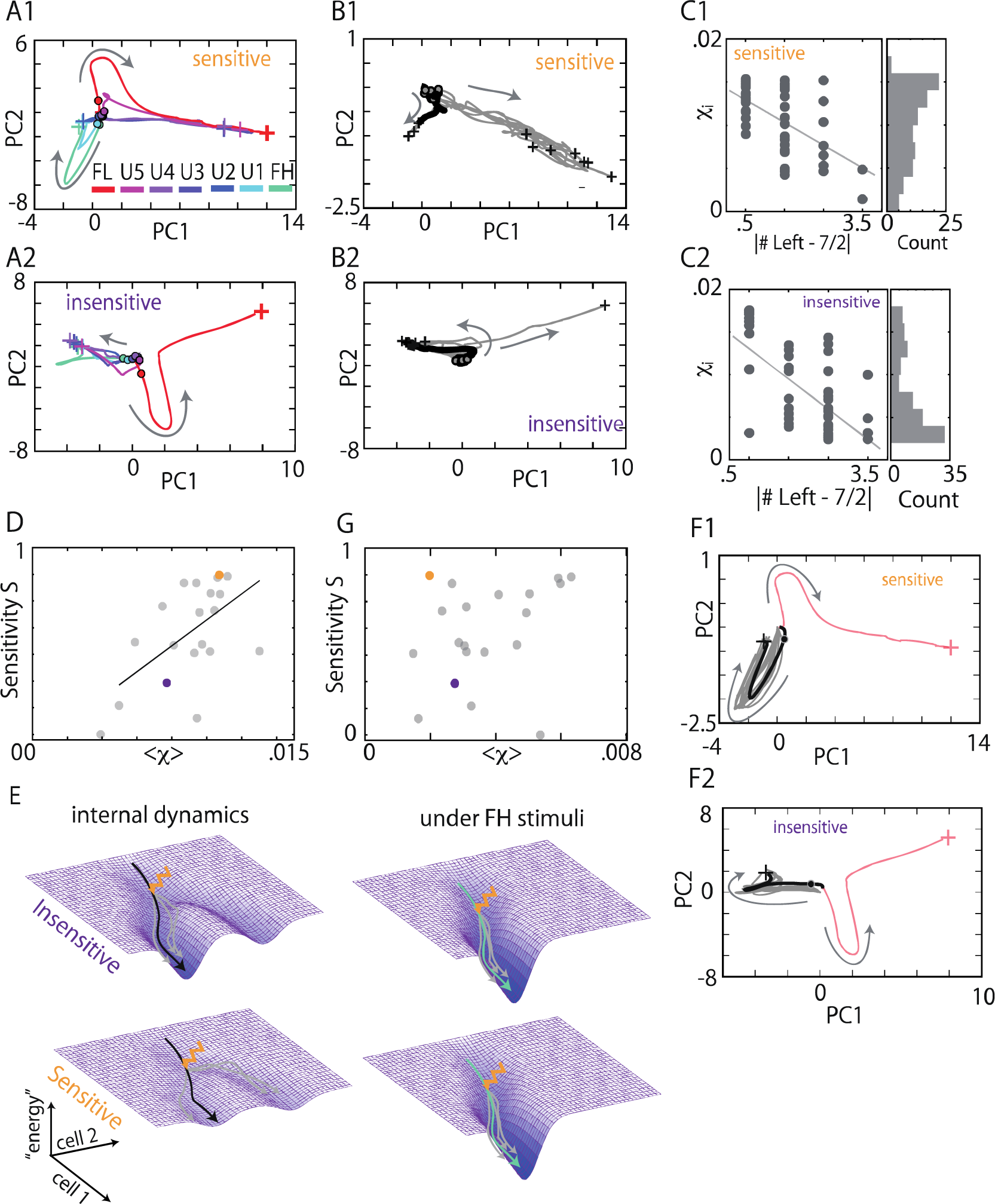
Characterization of neural dynamics for different choice behaviors. A1 and A2, Trajectories evoked by familiar and unfamiliar stimuli are shown in the two-dimensional PC spaces of sensitive or insensitive networks, respectively. B1 and B2, Ten perturbed trajectories (gray) and an unperturbed trajectory (black) in the absence of external stimuli are shown in the same networks and the same PC spaces as in A1 and A2, respectively. Gray circles refer to perturbed points and black crosses to decision points. All perturbed points are identical although they seemingly look different due to a temporal smoothing by a Gaussian function with the variance of 30 ms. C, Susceptibility *χ* (Methods) is shown for the same networks for stimulus-evoked trajectories evolving from 100 initial points of perturbation (left) against the response bias |#Left-3.5| (see the main text). Histograms of susceptibility are shown (right). D, Susceptibility and sensitivity are plotted for all successful learners. Susceptibility of each network was averaged over 50 initial points. E, Schematic images of the landscape of neural dynamics with (right) and without (left) familiar stimulus FH. F, Ten trajectories evoked by FH are perturbed at certain points (circles). Non-perturbed trajectories evolving from the same points are shown for FH (cyan). The non-perturbed trajectory evoked by FL is plotted in red for reference. G, Average susceptibility and sensitivity are plotted for all successful learners as in D. Susceptibility was calculated from the perturbed trajectories shown in E for FH.

We then examined a quantitative relationship between the susceptibility and responses to the seven stimuli (FL, U1, …, U5, FH). To this end, we define the response bias as |#Left - 7/2| for each perturbed state, where #Left is the number of Left choices resulting from each perturbed state in response to the seven stimuli. The response bias is 0.5 if three or four of seven choices are Left (weak biases), while it is 3.5 if all choices are only Left or Right (strong biases). Both in a sensitive (Figure 5C1) and an insensitive successful learner (Figure 5C2), the response bias is significantly and negatively correlated with the value of *χ*(p<10^−5^ for the sensitive and p<10^−6^ for the insensitive). We quantified the susceptibility of each model network by averaging the values of *χ* over randomly chosen 50 perturbed states. A network with a sensitive psychometric curve yielded a distribution biased towards higher values of *χ* (Figure 5C1), whereas a network with an insensitive psychometric curve yielded a distribution biased towards lower values (Figure 5C2). We plotted the sensitivity values of 20 successful learners against their average *χ* values to find a significant positive correlation (p<0.004) between the sensitivity (behavioral metric) and susceptibility (neural metric) (Figure 5D). These results are summarized as the following intuitive picture of the neural mechanism of individual differences: higher susceptibility implies a shallower landscape around the trajectory and produces a sensitive choice behavior, while lower susceptibility implies a deeper landscape and generates an insensitive choice behavior (Figure 5E).

In contrast, we did not find significant correlations between the sensitivity and the average susceptibility in the presence of learned stimuli (Figure 5G). Perturbed and unperturbed trajectories did not show large quantitative and qualitative differences in the presence of FH both in a sensitive and an insensitive network (Figure 5F). Similar results were obtained for FL (data not shown). Thus, our model suggests that internal network dynamics is an influential factor on the behavioral variability, but the influences are easily masked by external input (Figure 5E). This explains why we could not uncover a convincing relationship between neural population responses and behavioral variability in the rats.

### Trail-by-trial variance probes susceptibility

Susceptibility cannot be computed from the experimental data in rats. To find an alternative quantity to characterize the individual differences, we investigated trial-by-trial variances evoked by all stimuli in the models and rats, which was suggested to reflect the neural dynamics of decision-making [24, 25]. First, we calculated trial-by-trial variability before and after stimulus onset in the models (Methods). To reduce an artifact from neuron sampling, we normalized the post-stimulus-onset variability by the pre-stimulus-onset variability. Interestingly, the normalized variability is significantly correlated with the susceptibility in the models (Figure 6A), implying that trial-by-trial variance can be used as a proxy of the susceptibility. We also confirmed that the variability is correlated with the sensitivity in the models (Figure 6B1).

**Figure 6:**
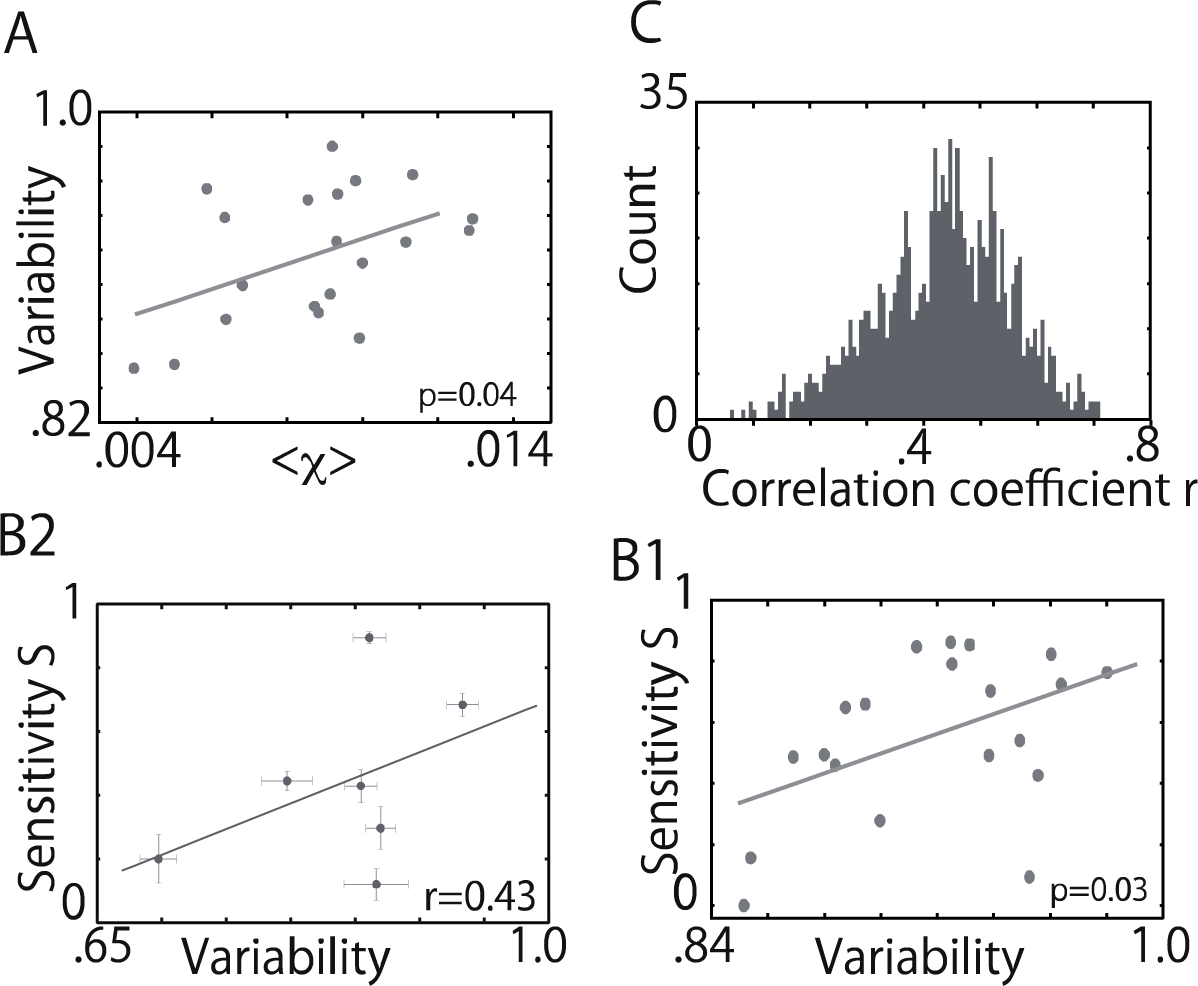
A, The susceptibility and the normalized trial-by-trial variability are significantly correlated in the all successful learners (p<0.04). B, Correlations between the normalized variability and the sensitivity across twenty successful learners and seven rats in B1 and B2, respectively. Data of one rat is excluded due to small number of available neurons (Methods). C, Correlation coefficients between the normalized variability and the sensitivity in rats. We resampled the experimental data and calculated 1000 the correlation coefficients for these data sets and plot them as a histogram.

Next, we calculated the normalized variability in the eight rats and examined its correlation with the sensitivity. (Figure 6B2). Due to the small number of the rats, we resampled experimental data 1000 times (Methods) and obtained 1000 resampled correlation coefficients between the normalized variability and sensitivity across the rats. All of the coefficients were positive (Figure 6C), indicating that neural population dynamics also reflects the behavioral differences of the individual rats and the normalized variability probes the characteristics (presumably the susceptibility) of neural dynamics.

### Reaction time is uncorrected with behavioral characteristics

RT is an important measure for the behavioral characteristics of individual animals. Next, we analyzed whether the sensitivity of psychometric curves is correlated with the RT of individual rats and models. The medians of RTs calculated for the eight rats were not significantly correlated with the selectivity (Figure 3S). These results show that the RT of rats does not strongly reflect the individual differences in choice behavior. We asked similar questions in our models and found that the RTs were also uncorrelated with the sensitivity of choice behavior in all the three cases, as in the rats.

However, we also noticed an interesting difference between the models and experiment. If we compare choice responses to familiar and unfamiliar cues in each model, RTs to unfamiliar cues were significantly longer than those to familiar cues (Figure S3D, p < 0.001). The result is consistent with our intuition. Somewhat unexpectedly, however, this was the case only in four rats, and even in these rats, differences in the median RT were typically as short as 10 to 20 ms (Figure S3H, p=0.8). The above discrepancy suggests that additional mechanisms not modeled here work in MFC during decision making in unfamiliar sensory conditions. We will argue the possible mechanisms in the discussion.

### Network structure only weakly reflects behavioral characteristics

Intuitively, network structure should be highly influential on neural dynamics and consequently on the individual behavioral differences. Therefore, we asked if network structure explains the behavioral characteristics and, if so, what aspect of the structure is relevant to them. Note that differences between the individual networks are considered to be small because they were obtained from similar initial networks (having the same weight distribution of recurrent connections) by training. In our statistical analysis of synaptic connections, crucial differences in network structure for producing significant differences in choice responses were not easily detectable. Actually, we tested various statistical indices to reveal a signature of such structural differences (Figure S4). Most of our attempts failed to find significant correlations between the statistical properties of network structure and the sensitivity. We, however, found an interesting exception when we analyzed a small portion of readout connections from such presynaptic reservoir neurons as exhibited the highest firing rates (Figure S4C). Thus, our results suggest that differences in network structure responsible for different behavioral characteristics do exist but they are subtle. Only a small fraction of connections that are actually used can be crucial for the observed behavioral differences.

### Discussion

We have shown how neural population in MFC processes a sensory-guided alternative decision making when rats are exposed to unfamiliar sensory stimuli. The choice responses to unfamiliar stimuli fluctuated over different rats and the psychometric curves of probabilistic choices largely varied between highly sensitive and poorly sensitive types across the individual rats. By using multi-electrode recordings, we have recorded spike trains of multiple medial frontal neurons during the task. We have built a recurrent network model and trained it through a reinforcement learning algorithm to successfully reproduce both temporal evolution of neural population activity and spectrum of observed choice behaviors. By studying neural dynamics in this model, we have found that the susceptibility of internal neural dynamics predicts individual differences across different networks. This modeling result predicts that highly sensitive rats should exhibit a large trial-by-trial variability in neural trajectories, which was also confirmed in the rats.

### Reward-guided neural trajectory learning

Neural activity sequences have been ubiquitously found in the mammalian brain executing behavioral tasks including decision making [4, 5, 26, 27]. In the rat MFC, neural responses to different auditory stimuli subsequently converged to choice-specific trajectories [9]. This study has further demonstrated the relevance of the dynamical characteristics of choice-specific neural trajectories in MFC to the diverge spectrum of animals’ choice behaviors.

To explore the neural mechanisms of ambiguous decision making for unfamiliar stimuli, we have modeled a reservoir network [11–14] that undergoes reinforcement learning. By employing a eligibility trace [15–17], we trained the network model to find the formation of familiar trajectories in the reservoir which eventually converge to one of the two local areas corresponding to choice responses in the state space. Neural trajectories driven by unfamiliar stimuli stochastically diverged into the vicinities of these decision areas organized for familiar stimuli with a stimulus-dependent probability, resulting in ambiguous choice responses.

Separation of neural trajectories occurs similarly in the rats and models. However, the separation somewhat slows down in the rats about 100 ms after the cue onset during the cue presentation (Figure 3E), whereas the separation continues in our model without slow down (Figure 3F). Indeed, around this timing a sharp increase in the gamma power of the local field potentials occurred in the MFC [9]. As suggested in the rodent [28, 29] and primate [30], the increased gamma oscillation may indicate the onset of cross-area communications in the MFC, presumably for the preparation of behavioral outputs. This communication across areas, which is not implemented in our model, might suppress the separation.

### Correlations between neural characteristics and behavioral variability

The behavioral variability of individuals have been a focus of psychological and behavioral studies of humans [31]. Choice preferences of individuals under uncertain or risky conditions have been studied in the context of value evaluation by a large-scale brain network including cortical and subcortical structures [32, 33]. Furthermore, recent studies using fMRI have revealed correlations between task performance and intercortical functional connectivity, indicating that macroscopic brain activity influences the behavioral variability of individual animals [34–36]. However, relatively little has been known about the neural substrates for the individual differences [37]. Our results suggest that neural dynamics in a local cortical area (of MFC) is a determinant of individual differences in the rapid choice behavior that does not require strategic exploratory decisions. We have shown evidence for a relationship between sensitivity in psychometric curves and the dynamical properties of neural trajectories (such as susceptibility) formed during sensory experience (Figure5D).

Recent studies have revealed that neural processing is not a simple passive filter, but an active dynamical process [26, 27, 38–40]. From the dynamical system viewpoint, the flow structure of neural state space, such as line attractors [26], separatrix [27] and “null space,” i.e., the subspace of neural state space of motor cortex on which neural trajectories are uncorrelated with movements, but influence the choice of subsequent movement [40], has a significant influence on linking externally or internally driven information to behavioral output.

In addition to these structures, our model suggests crucial influences of covert properties of intrinsic neural dynamics, such as the susceptibility, on decision making in an unfamiliar sensory environment. We demonstrated that the susceptibility of neural dynamics to perturbation in the absence of sensory stimuli predicts the sensitivity in individual psychometric curves (Figure 5D). If the susceptibility is high in a model, psychometric curve of the model tends to be sensitive to differences between unfamiliar stimuli. Recent studies have revealed neural mechanisms of sensory information coding [41], short-term memory [42], and motor planning [43] by applying perturbation to neural activity with opt-genetic techniques. Our results suggest that response to perturbation reflects a part of individual difference. Our model also gives a testable prediction that animals in which neural activities are strongly disturbed by optical stimuli to MFC have highly sensitive psychometric curves in response to unfamiliar stimuli.

Correlations between the susceptibility in neural dynamics and the sensitivity in behavior were not obvious when neural dynamics was driven by familiar stimuli (Figures 5F and 5G). This implies that behaviorally relevant differences in intrinsic neural dynamics are not necessarily observed under the learned conditions. This might be natural when all subjects are trained to perform the same task. Previous computational models [44–46] and experimental studies [36] demonstrated the impacts of intrinsic neural dynamics on processing external stimuli or cognitive tasks. Our results suggest that intrinsic neural dynamics can influence the characteristics of individual behavioral responses.

We found that the susceptibility predicts large trial-by-trial variations in neural activity in our model and that trial-by-trial variation is in turn correlated with the individual differences in both models and rats (Figure 6). In humans, larger trial-by-trial variability in movements predicted faster rates of motor learning, though whether high variability in movements implies high variability in neural activity remains unclear [47, 48]. In the present task, large variability in neural responses may increase the flexibility in controlling neural trajectory evolution, thus enabling the rats to flexibly adjust their probabilistic behavioral responses according to cue tones. This may explain the observed correlations between the trial-to-trial variability and psychometric curves. However, the correlations were weak in the rats (Figure 6), and this point requires further experimental clarifications.

### Network structure weakly indicates the sensitivity in psychometric curves

We have investigated the relationships between several metrics of network structure and the sensitivity in psychometric curves of the networks to find no significant correlations in most of the cases. We have only found that the Left-Right asymmetry in the weight sums of readout connections from the most active reservoir neurons indicate the sensitivity of different network models (Figure S4, p<0.03). However, neither readout connections from all reservoir neurons nor the sums of the strongest connections were significantly correlated with the individual differences. All together, these results suggest that network structure alone cannot unambiguously distinguish the sensitivity.

Our learning procedure modifies only readout connections. Though this assumption is unrealistic, our model could form the neural trajectories that successfully associate sensory inputs to correct behavioral responses. Thus, our results suggest that learning a simple association like the present task can be completed through modifications of inter-cortical-area connections or cortico-subcortical connections. However, learning a complex task may require modifications of intra-cortical-area connections for organizing an adequate neural network.

A log-normal distribution of synaptic connections, which is adopted in our model, has been found in local cortical circuits [22, 23]. Some studies [21, 23] have suggested that this class of connection is helpful to generate a rich variety of spike sequences in spontaneous activity. Therefore, the log-normal connection in the reservoir likely contributes to generating neural trajectories specific to stimulus and choice (Figures 3A and 3B) and, consequently improves learning performance. A role of the log-normal connection on learning is left for a future work.

### Different RT profiles between rats and models

We may speculate that rats with highly sensitive psychometric curves carefully evaluate sensory information before making behavioral responses and hence tend to require a longer time for decision making. However, this naive speculation was not supported by our experimental results [9]. Actually, we have found in both rats and models that RTs are uncorrelated with the sensitivity for both familiar and unfamiliar cues. The present results for RTs are largely consistent between the rats and models, but we have found a few exceptions. All of the rats made rapid decisions even for unfamiliar cues (Handa et al., 2017), yielding comparable RTs for familiar and unfamiliar cues (Figure S3C: the mean RTs differed by at most 20 to 30 ms between the two cue types), whereas in our model the mean RT for familiar stimuli was much shorter than that for unfamiliar ones although these differences were also uncorrelated with the sensitivity of the models (Figure S3G).

How this discrepancy arises remains unclear, and we only speculate the possible underlying mechanisms. The discrepancy seems to indicate that the MFC recruits additional mechanisms of decision making, which was not incorporated into the present models, when the rats are exposed to unfamiliar cues. A likely explanation is that threshold for decision making is increased to improve the accuracy of behavioral responses to familiar cues through the learning of sensory environment. Yet another explanation is that unfamiliar bottom-up signals from primary sensory cortices to MFC is less effective in activating medial frontal neurons, as they have not learned the unfamiliar inputs. Indeed, modeling studies suggest various plasticity mechanisms to gate the learned synaptic inputs effectively and robustly [49, 50]. The causes of the discrepancy are open to future studies.

## Acknowledgements

We thank Joshua Johansen for critical comments on the manuscript and Masami Tatsuno for useful discussion about analysis of correlation. This work was partly supported by KAKENHI (nos. 16H01289 and 17H06036 to T.F.) from MEXT.

## Methods

### Experimental procedure

All experiments were carried out according to the Animal Experiment Plan approved by the Animal Experiment Committee of RIKEN. The details of experimental procedure are given in [9]. Briefly, head-restrained adult Long-Evans rats (male 210-240 g: SLC) were trained to associate licking of spouts with reward delivery. We presented either of two pure tones (*familiar* cue tones: 13.0 and 10.0 kHz, for 0.2 s) in a pseudo-random order as a cue for licking a spout located at the left or right side of rat, respectively (Figure 1A). The rats were required to lick a correct spout within 5.0 s from the cue onset to obtain a reward (0.1% saccharin water). If the licking response was incorrect, we did not deliver the reward and prolonged the duration of the immediately following post-response period (3.0 s) by 5.0 s as an aversive experience. We continued the training until the correct rate finally reached a criterion (75%) without error correction. Each rat underwent one or two days of subsequent recording sessions, in which we presented the *two familiar* cue tones (10.0 and 13.0 kHz) and five *unfamiliar* cue tones (10.5, 11.0, 11.5, 12.0 and 12.5 kHz) with the occurrence probability of 80% or 20% (4% for each unfamiliar tone), respectively. The correct familiar cue trials were always rewarded, whereas the reward probability was linearly varied for unfamiliar cue trials along its cue tone frequency: (Left/Right) = 10.5-kHz (0.17/0.83), 11.0-kHz (0.33/0.67), 11.5-kHz (0.5/0.5), 12.0-kHz (0.67/0.33), 12.5-kHz (0.83/0.17). We trained 36 rats with familiar tones and only 21 reached the criteria for successful learning. After surgery, 15 were available for multi-neuron recordings and eight of them finally yielded qualitatively and quantitatively satisfactory data for the succeeding analysis.

We recorded multiunit activity mainly from the deep layers (depth from pia matter: 1.0-2.0 mm) of the MFC (+2.7-+3.6 mm anterior, 0.6-2.0 mm lateral of Bregma) through a 32-channel silicon probe consisting of 4 shanks (Neuro Nexus Technologies, Inc., USA), each with 2 tetrode sites separated vertically by 0.5 mm. We only analyzed the behavioral and neuronal data obtained on the first day of the recording sessions when the rats were still not habituated to unfamiliar cues. Spikes were sorted with a custom-made semi-automatic spike sorting program, EToS [51], and the sorted spike clusters were further analyzed manually using Klusters and NeuroScope [52, 53]. The number of isolated units (RS + FS) was n=37 in rat #941, n=68 in #897, n=22 in #902, n=12 in #807, n=44 in #880, n=40 in #879, n=50 in #940 and n=51 in #949. Neuronal activity and behavioral performance were analyzed using MATLAB (The MathWorks, Inc.).

### Neuron models

We constructed a network model composing three parts, an input layer, a reservoir network and a readout layer (Figure 2A). The input layer has 200 neurons, each of which responds to stimulus *k* at the following firing rate *r*_i_:

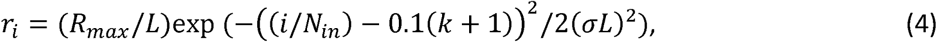

where *R*_max_, *N*_in_, and *σ* are the maximum firing rate, the number of input neurons, and the standard deviation, respectively. The values of these parameters were set as *R*_*max*_ = 110, *N_in_* = 200 and *σ* = 0.15, and *k* (= 1, …, 7) specifies the preferred stimulus of the neuron, with *k* = 1 and 7 corresponding to familiar high (FH) and low (FL) stimuli, respectively, and *k* = 2, …, 6 to five unfamiliar inputs (U1-5), respectively. The parameter *L* is the width of frequency tuning curves, and most results were calculated for *L* = 0.4 except in Figure 3G.

The reservoir network is essentially the same as the recurrent network model studied in Teramae et al. [21] except for the introduction of NMDA receptors and background noise as well as minor modifications of model parameters. The reservoir network has 5000 excitatory and 1000 inhibitory leaky integrate-and-fire neurons obeying the membrane dynamics

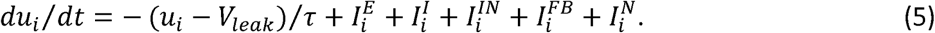

where *V*_leak_ = −70 [mV], *τ*= 20 or 10 [ms] for excitatory and inhibitory neurons, respectively, and the refractory period is 1 [ms]. If *u* reaches threshold V_th_ = −45 [mV], the neuron fires and *u* is reset at *V*_r_ = −60 [mV], Excitatory and inhibitory recurrent synaptic inputs, 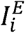 and 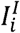, are described by the following equations:

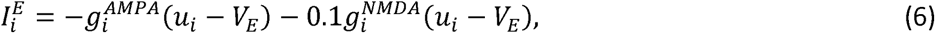

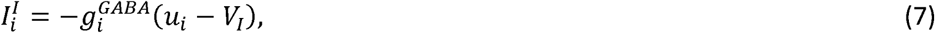

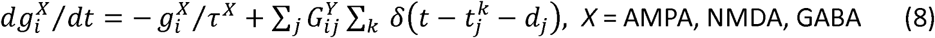

where 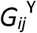 and *d_j_* are the weight and delay of synaptic connection from neuron *j* to neuron *i*, respectively, and 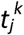 is the *k-th* spike time of neuron *j.* The reversal potentials of synaptic inputs are *V*^E^ = 0 [mV] and *V*^I^ = −80 [mV], synaptic time constants are *τ*^AMPA^ = 8 [ms], *τ*^NMDA^ = 100 [ms] and *τ*^GABA^ = 8 [ms], and δ is the Kronecker’s delta function. All synaptic conductances *g* are normalized by the membrane capacitance to have the dimension of [1/ms]. At excitatory-to-excitatory (E-to-E) connections, synaptic delays are chosen randomly from 1 to 3 [ms] and at other connections they are from 0.5 to 1.5 [ms]. Synaptic input from input neurons 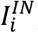 feedback input 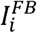 from readout neurons (see below), and background noise 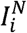 are all excitatory and defined as

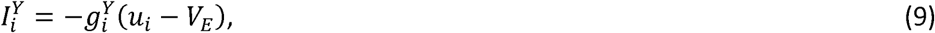

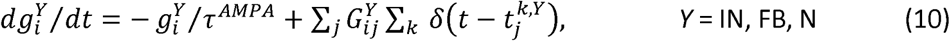

where 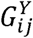 is the synaptic weight from neuron *j* to neuron *i* and 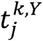 is the *k*-th spike time of presynaptic neuron *j* of input type *Y*. In the present study, *G*^N^ = 0.1, and other weights obey lognormal distributions, as described later. In Equation 10, spikes in *I*^IN^ and *I*^FB^ are generated by non-stationary Poisson processes with the instantaneous firing rates given by Equation 4 or those of the readout neurons, respectively. The feedback spikes are generated independently for individual postsynaptic reservoir neurons to avoid strongly correlated activation of these neurons. Background noise is given by a Poisson spike train of 20 [Hz].

The readout layer has two rate-based neurons, L- and, R- neuron, which integrate spike inputs from the reservoir and undergo mutual inhibition. Their firing rates *r_L,R_* obey

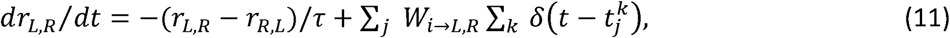

where *τ* = 50 [ms]. The value of *r*_*L,R*_ is set to zero if it takes a negative value due to the mutual inhibition.

### Network architecture

Among 5000 excitatory neurons in the reservoir network, 2000 neurons receive excitatory external input from input neurons through sparse connections. The connection probability *p*=0.01 and their weights are set at 0.02. The wiring probabilities of excitatory-to-excitatory (E-to-E), excitatory-to-inhibitory (E-to-I), inhibitory-to-excitatory (I-to-E) and inhibitory-to-inhibitory (I-to-I) connections are *p*^EE^ =*p*^EI^ =0.1 and *p*^IE^ =p^II^ =0.5. The weights of E-to-E connections are generated according to the following log-normal distribution:

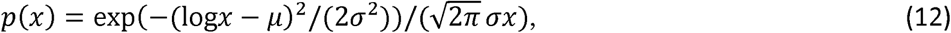

where σ^EE^ = 0.8 and μ^EE^ = log(0.01)+(σ^EE^)^2^. The weights of E-to-I, I-to-E and E-to-I connections are fixed at 0.01, 0.002 and 0.0025, respectively.

Each readout neuron is projected to by a subset of reservoir neurons not receiving sensory input through readout connections *W*_i->L,R_ and feed their outputs back to another subset of reservoir neurons. The former and latter subsets of neurons consist of randomly and independently chosen 30% and 50%, respectively, of such reservoir neurons as are connected to none of the input neurons. An overlap can exist between the L-neuron-projecting and R-neuron-projecting subsets. The weights of *W*_i->L,R_ are randomly chosen between 0 and 60. The weights of feedback connections *G*^FB^ obey a lognormal distribution given in Equation 12, with the variance and mean given as *σ*^FB^ =1.2 and *μ*^FB^=log(0.012)+(*σ*^FB^)^2^.

### Learning protocol

The present network model learns to correctly associate external stimuli with choice responses by modifying the readout connections *W*_i->L,R_ through reinforcement learning. The learning procedure is as follows: first, the network undergoes a pre-stimulus run for 100 [ms], which induces a baseline activity in the initial network state; subsequently, one of the familiar stimuli (FH or FL) is applied for 200 [ms]; after the termination of stimulus followed by a delay period of 50 [ms], the network is allowed to make a decision if the activity of a readout neuron exceeds that of the other readout neuron by a criterion difference *θ* (Figure 3B). If the network does not reach a decision criterion by 900 [ms] from the end of delay period, the trial is reset and a novel trial is initiated. If the decision by the network is correct, the readout weights are modified with a positive reward, whereas the weights are punished with a negative reward if the decision is incorrect. If the network fails to give a decision in a trail, the network is also punished. If the network reaches a decision criterion before the end of delay period, we discard this trial and start a new trial.

Readout connections are modified in terms of eligibility trace *e*_i->L,R_ according to Equations 1-3 [15–17], The eligibility trace is assigned to each readout connection and measures the extent to which a particular connection contributes to decision making. The variables *a*_*i*_(t) and 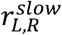 in Equation 1 are computed in terms of presynaptic and postsynaptic activities as

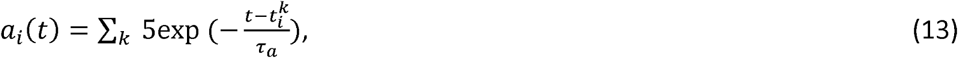

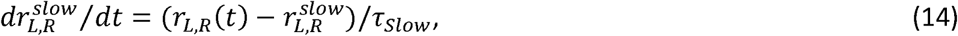

where *τ*_*a*_ =10 [ms] and *τ*_*slow*_ = 50 [ms]. The term 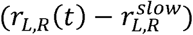 detects a rapid change in *r*_*LR*_(*t*) faster than *τ*_*slow*_ so the eligibility trace is increased by coincidence between high presynaptic firing rate and a rapid increase in the readout activity. In modifying readout connections in Equation 3, we use a normalized eligibility trace defined as

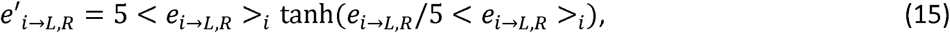

where < … >_*i*_ denotes averaging over presynaptic reservoir neurons. We modify readout connections by reward expectancy *U* and the limited eligibility according to Equation 3. We manually bound the value of *W*_i->L,R_ (T) below 60. Reward expectancy and decision criterion are adaptively modified during learning as follows:

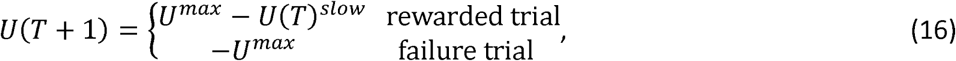

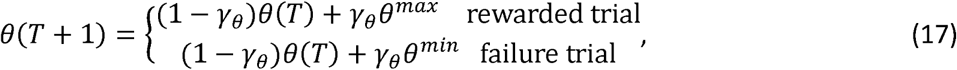

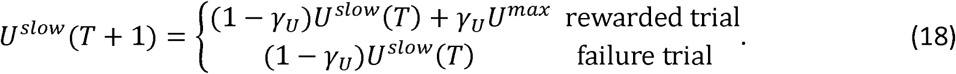

At an arbitrary *T*, −*U*^*max*^ ≤ *U*(*T*) ≤ *U*^*max*^ and *θ^min^* ≤ *θ* ≤ *θ*^*max*^. Parameter values are *U^max^* = 1, *θ*^*min*^= 10[Hz], *θ*^*max*^= 30[Hz], *γ*_*U*_ = 0.05 and *γ*_*ϑ*_ = 0.005. The variable *U*^slow^(*T*) is rapidly decreased while the criterion θ(*T*) is increased when the network has been rewarded in successive trials. Therefore, we may regard *U^slow^*(*T*) as reward expectancy at the learning step.

### Fitting psychometric curves

Psychometric curves were calculated from the probability of Left choices. In the training of our model, the criterion of decision *θ* was initially kept low and then gradually increased until it was finally fixed at 50 [Hz] after learning. This manipulation made the separation of neural trajectories easier and clearer at the decision timing without changing the qualitative behavior of the model with constant *θ*. The Left-choice probabilities for familiar and unfamiliar stimuli were calculated for each rat over a few hundred or a few tens of trials, respectively. The probabilities for all stimuli were calculated for each network model over 50 trials except in Figures 5C in which we simulated 100 trials. In a small fraction of trials (< 1%), the network model did not reach the decision criterion within the time limit for simulations (< 900 ms). For such trials, we assigned a “relative distance” to the decision criterion *θ* at the time limit to the Left-choice probability: 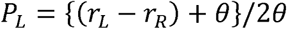.

Unless otherwise stated, we fitted psychometric curves (the probability of Left choices) by a nonlinear function in the least square method: *P*_L_ = *c*(tan(*f* - *a*) + *b*). The variable *f* presents a normalized tone frequency {−1, −2/3, −1/3, 0, 1/3, 2/3, 1}, which corresponds to {10, 10.5, 11, 11.5, 12, 12.5, 13} [kHz] in the rats, respectively, and {FL, U5, U4, U3, U2. U1, FH} in the models, respectively. After fitting a psychometric curve, we normalized the parameter *c* to define the sensitivity *S* such that *S* = 1 refers to a linearly increasing psychometric curve *P*_L_(*f*) =(*f*+1)/2 and S = 0 to a highly biased psychometric curve, *P*_L_(−l)=*P*_L_(−2/3)= … =*P*_L_(+2/3)=0 and *P*_L_(1)=1. The larger the value of *S*, the more sensitive a psychometric curve. A conventional sigmoid function was not suitable to characterize sensitivity in psychometric curves.

We examined whether the evaluation of sensitivity remains stable for each rat and model if a different fitting scheme was used to characterize their psychometric curves. In this scheme, we used *Pʹ(f) = bʹ* tan(aʹ*f*) + (1 - *b*ʹ tan(*a*ʹ)) as a new fitting function, and defined a parameter as *s*ʹ = *a*ʹ × *b*ʹ (Figure S1B-E). This definition is reasonable as *a*ʹ and *b*ʹ scale the x-axis and y-axis, respectively, to modify the slope of the fitting function. Then, a new sensitivity *S*ʹ was defined from *s*ʹ by the normalization process mentioned previously.

### Linear discriminant analysis

We examined the neural dynamics underlying decision making in the rats and models by Linear (Fisher) discriminant analysis (FDA). We grouped neural states of familiar trajectories at the time of decision making (in the rats, the time of licking responses) into two groups: a group of neural states in Left-choice trials and a group of neural states in Right-choice trials. FDA identifies such an (*N*-1)-dimensional hyperplane that maximizes the ratio of the mean distance between the two groups (inter-group distance) to the sum of standard deviation from this hyperplane in each group (intra-group distance). Here, *N* is the dimension of neural state, i.e., the number of neurons in the population. We defined a one-dimensional line, called *W*opt, that is orthogonal to the identified (*N*-1)-dimensional hyperplane.

### Linear regression analysis

In addition to FDA, we used a regression method to identify the two (choice and stimulus) axes explaining differences in trajectories between the Left and Right choice conditions in the rats and between FH and FL stimuli in the models. Our method is the same as used in [26, 27]. Briefly, we obtained the average firing rates of 100 most active neurons across trials under given stimulus and choice condition in the model and the average rate of 30 neurons which showed the largest differences in firing rate between pre- and post-stimulus onset in the rats. We used neural activities recorded from 100 (100) ms before to 500 (300) ms after the stimulus onset in models (rats). After gaussian-filtering, we calculated the z-scores of these activities, where the standard deviation of the filter was 10 ms for the models and 30 ms for the rats. Then, we performed a linear regression analysis,

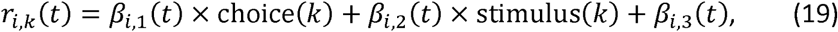

where *r_i,k_*(*t*) is the z-scored responses of neurons (*i* = 1 - 100) at time *t* on trial *k*, and *β*_*i*,1_(*t*), *β*_*i*,2_(*t*) and *β*_*i*,3_(*t*) represents the choice coefficient, stimulus coefficient and residual component, respectively. Here, these coefficients are projected onto the subspace spanned by the ten largest PCs, and choice (*k*) and stimulus (*k*) are binary variables.

### Susceptibility

Susceptibility *χ_u_* characterizes the dynamical trends of a neural network by measuring how it evolves in response to a perturbative input given to neural state *u*. To measure *χ_u_*, we set the membrane potentials in all reservoir neurons at the initial value of −60 [mV] and simulated the time evolution of neural population up to time *t_0_* (=100 [ms]), at which we applied a perturbative input to the trajectory: 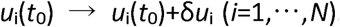. Choosing a different direction of perturbation ***δu*** randomly in every trial, we obtained 30 different perturbed trajectories ***u***^(*k*)^ (*k* = 1,…,30) all of which were perturbed at the same state {*u*_i_}. Each component of ***δu*** is chosen from a uniform probability distribution of [−0.1, 0.1] [mV]. To evaluate the effect of the perturbation on state evolution, we evaluated the average distance *D* between the perturbed trajectories and an unperturbed trajectory ***u***^(0)^ at time *t*_1_ = 500 [ms] as follows. Let 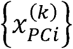 be the *i*-th PC of neural state at time *t_1_* along the *k-*th perturbed trajectory ***u***^(*k*)^ (*t*_1_), where *k* = 0 refers to the unperturbed trajectory for convenience. If the network model reaches the decision criterion before time 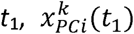 is calculated at this time. We define *χ_u_* for the perturbed state *u* as

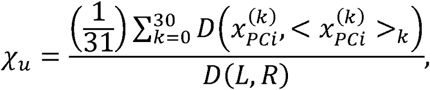

where 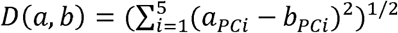 is the Euclidean distance between state *a* and state *b* on the five-dimensional PC subspace, 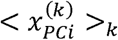 is the average of the perturbed and unperturbed trajectories, and the distance from the average trajectory is averaged over the trajectories. *D*(L, R) provides a normalization factor, and is the distance between Left-choice and Right-choice points on the PC subspace, where the choice points refer to the averaged neural states at which the network model reaches Left or Right choice.

We emphasize that the susceptibility is assigned to each neural state rather than each trajectory. Therefore, a neural trajectory could show a broad range of the susceptibility depending on the specific state at which a perturbative input was given. In Figures 5A,B,and F, we plotted perturbed neural trajectories starting from various neural states with low to high *χ*_*u*_ values. In Figures 5D and G, we defined the susceptibility of a model network by summing up the *χ*_*u*_ values of 50 neural states.

### Trial-by-trial variability in rats and models

We selected those neurons that exhibited average firing rates greater than 3Hz during the whole task period, and applied PCA to the activity of this neural ensemble during a pre-stimulus epoch (a 100 ms-long interval prior to stimulus onset) and a post-stimulus epoch (a 300 ms-long interval following stimulus onset). We computed the logarithm of cumulated product of PC variances, 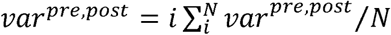, where *N* is the number of selected neurons and *vari^pre,post^* is the variance of the *i*-th PC for pre- and post-stimulus ensemble. We performed the analysis in seven rats and 20 successful learners. One rat was excluded as it yielded only one neuron according to the aforementioned criterion. Note that, in applying PCA, we randomly picked up 30 trials for each stimulus and each rat because the number of recording trails varies from rat to rat and from stimulus to stimulus. We normalized the post-stimulus variance by dividing the pre-stimulus variance to define the normalized variability. In Figure 6C, we calculated the sensitivity *S* for the resampled trials, and then calculated correlation coefficients between the resampled variabilities and resampled sensitivities in the seven rats.

